# Profiling the impact of the promoters on CRISPR-Cas12a system in human cells

**DOI:** 10.1101/2023.02.06.527393

**Authors:** Jinhe Li, Qinchun Liang, HuaPing Zhou, Ming Zhou, Hongxin Huang

## Abstract

The plasmid vector platform is the most commonly used for the expression of the versatile CRISPR-Cas technique and the promoter is a crucial element for the expression vector, thus profiling the impact of the promoters on CRISPR editors provides the basic information for the gene-editing toolkits and can be a guideline for its design. Herein, we made a parallel comparison among four commonly used promoters (CAG,∼1700bp; EF1a core, ∼210bp; CMV, ∼500bp; and PGK, ∼500bp) in CRISPR-Cas12a system in mammalian cells to explore the impact of promoters on this powerful tool. We found that without badly damaging targeting specificity, the CAG promoter-driving Cas12a editor exhibited the most active (efficiency takes as 100%, specificity index= ∼75%) in genomic cleavage, multiplex editing, transcriptional activation, and base editing, followed by promoter CMV (efficiency=70∼90% (vs CAG), specificity index= ∼78%), and then EF1a core and PGK (both efficiency=40∼60%, vs CAG) but with higher specificity (specificity index= ∼84% and ∼82%, respectively). Therefore, CAG is recommended in the CRISPR-Cas12a system for the applications that need a robust editing activity but without size limitation, CMV mostly can be an alternative for CAG when requiring a smaller space, EF1a is similar to PGK with relatively high specificity, but has a smaller size, thus is more suitable for *in vivo* therapeutic applications. The data outlined the properties of the widely used promoters in the CRISPR-Cas12a system, which can be a guide for its applications and can be a useful resource for the gene-editing field.

## Background

The CRISPR-Cas systems have been harnessed as powerful tools for a variety of basic research and clinical therapy, including programmable genome editing[1, 2], gene activation[3-5], live imaging[6, 7], base editing[8, 9], and primer editing[10, 11]. However, the successful application of this versatile technology requires the essential expression of the Cas-nuclease protein, which can be generated by a rational synthetic design of the expression cassettes[12]. The plasmid vector platform is the most commonly used for the expression of the CRISPR-Cas tools, such as the adeno-associated virus (AAV) system, the most widely used viral vector for *in vivo* gene-editing tools’ delivery[13-15]. Plasmid vector systems contain cis-acting elements, such as enhancers, promoters, polyadenylation signals, and other expression elements, all of which can affect the expression levels of the transgene. To achieve high transgene expression levels, several investigations had been reported by optimizing cis-acting elements in plasmid systems[16-19]. For example, the enhancer/promoter is a critical element in an expression vector, and the selection of the CAG promoter had been reported to significantly increase the expression and the stability of the transgene in mammalian cells[16]. Although it had been shown that different promoters had different effects on the expression level of the interesting gene in the plasmid vector[16-20], the effects of the promoters on CRISPR gene-editing tools, such as editing activity, targeting specificity, transcriptional activation level, and base editing ability, have not been comprehensively elucidated. Reducing the amount of the Cas-protein in the cells improved the targeting specificity but affected the efficiency[21-23]. In addition, an *in vivo* therapy purpose application always required a more accurate editing[4, 13-15]. Therefore, there is a need for considering the level of the Cas-protein in CRISPR-Cas plasmid systems, and a rational design of a CRISPR editor vector can make this technique play a better function and can be applied to the fitness of their unique properties to the intended purpose.

In the current study, we analyzed the effects of four commonly used promoters (CAG, EF1a core, CMV, and PGK) on the CRISPR-Cas12a system, including cleaving activity, targeting specificity, multiplex editing efficiency, gene activation level, and base editing ability, which provided the basic information about the impact of promoters for the CRISPR gene-editing toolkit.

## Methods

### Plasmids construction

The plasmids expressing the Cas12a-nucleases used in this study were designed with a promoter (CAG, EF1a Core, CMV, PGK, for the detailed sequences see Supplementary information) driven enAsCas12a-HF CDS, 3xHA, and a P2A-mcherry reporter, or a promoter (CAG, EF1a Core, CMV, PGK) driven dAsCas12a-HF CDS fusing with VP64-p65-Rta (VPR) activation domain, or a promoter(CAG, EF1a Core, CMV, PGK) driven rat APOBEC1 fusing with dAsCas12a-HF CDS. All the plasmids’ cloning was constructed by standard PCR via Gibson Assembly. The sgRNA expression plasmids were constructed by ligating oligonucleotide duplexes into the backbone with a human U6 promoter and an AsCas12a-crRNA scaffold sequence. All the plasmids were confirmed by Sanger sequencing and all the sgRNAs oligonucleotides used in this study are shown in Supplementary information Table S1.

### Cell culture

HEK293T cells (ATCC, CRL-3216) were maintained in Dulbecco’s Modified Eagle’s Medium (DMEM, Life Technologies) and MCF7 cells (ATCC, HTB-22) were cultured in RPMI 1640 medium (Life Technologies) at 37°C in a 5% CO2 humidified incubator. All the media contained 100 U/mL penicillin, 100 μg/mL streptomycin (Life Technologies), and 10% fetal bovine serum.

### Cell transfection

For cell transfection, approximately 2.0×10^5^ cells were seeded in the 24-well plate, and the following day when cells grew up to ∼70%, the transfection was administered by the polyethyleneimine (PEI) method. To detect the expression level of the mCherry reporter or the Cas-protein, MCF7 or HEK293T cells were transfected with 100ng of Cas-nuclease expression plasmid per well in a 24-well plate. To detect the disruption of the mNeonGreen reporter, HEK293T-KI mNeonGreen cells were transfected with 120ng of Cas-nuclease expression plasmid and 80 ng of the sgRNA-encoding plasmid per well in a 24-well plate.

### Tag-seq experiment and analysis

Tag-seq experiments were used to compare and analyze the specificity of the CRISPR-Cas12a systems with different promoters, which were performed as previously described [24, 25]. Briefly, HEK293T and MCF7 cells were transfected by PEI with 10 pmol Tag, 600 ng of Cas nuclease, and 600 ng pool sgRNAs (25 guides) or a single crRNA array (targeted 6 sites) per well in a 12-well plate. Three days post-transfection genomic DNA was extracted for one-step libraries preparation by the Fragmentation, End Preparation, and dA-Tailing Module and Adapter Ligation Module kit (Vazyme Biotech Co., Ltd., Nanjing, China). The libraries were constructed by PCR with library preparation primers, and purified by Hieff NGS™ DNA Selection Beads (YEASEN, China), followed by sequencing (NovaSeq platform, Novogene, Beijing, China) and analyzed with a Tag-seq bioinformatics pipeline (https://github.com/zhoujj2013/Tag-seq).

### Deep-seq experiments and base-editing analysis

Deep-seq experiments were used to assess the base editing efficiency of the CRISPR-Cas12a systems with different promoters. HEK293T and MCF7 cells were transfected by the PEI method with 600 ng of Cas nuclease, and 400 ng sgRNAs per well in a 12-well plate. Two days post-transfection genomic DNA was extracted for deep-seq libraries preparation. Briefly, the primers were designed with forward and reverse indexes to amplify the genomic sequence in the first-round PCR. Then, an equal amount of the first PCR products with each sample was pooled and administrated to a second round of PCR with the primers containing the P5 and P7 motifs to generate a standard library. Paired-end sequencing was used by the NovaSeq platform (Novogene, Beijing, China). The base editing results were analyzed using the batch version of the CRISPResso2[26]. The deep-seq primers were listed in Supplementary information Table S2.

### Western blotting

To examine the expression of the Cas12a, MCF7 cells were transfected with the Cas12a encoding plasmids using the PEI method. Briefly, the transfected cells were collected 2 days post-transfection and then lysed in a 2×SDS loading buffer for boiling for 10 min. Lysates were resolved through SDS/PAGE and transferred onto a nitrocellulose membrane which was blocked using 5% non-fat milk and sequentially incubated with primary antibodies (anti-HA, sigma, USA; anti-GADPH, Proteintech, China) and an HRP-conjugated horse anti-mouse IgG secondary antibody (CST, USA). All the probed proteins were finally detected through chemiluminescence following the manufacturer’s instructions.

### FACS analysis

All the flow cytometry results were analyzed by FlowJo software. For detection of the transfection efficiency, the transfected cells were obtained 2 days post-transfection and then subjected to a flow cytometry by calculation of the proportion of the mCherry reporter. For detection of the disruption efficiency of the mNeonGreen reporter, HEK293T-KI mNeonGreen cells were harvested 2 days post-transfection and the editing efficiency was determined as the proportion of mNeonGreen negative cells within the Cas-nucleases transfected cells (mCherry-positive).

### Quantitative real-time PCR

The activation ability of the CRISPRa activators was determined by qPCR methods. Detailedly, the total RNA from the transfected cells was extracted by Trizol Reagent (Thermo Fisher, USA) approach according to the manufacturer’s instructions. 1 ug of the total RNA was reverse transcribed into cDNA and the quantitative real-time PCR was performed using a LightCycler 96 System (Roche, Switzerland). Relative gene expression was calculated using the 2^−ΔΔCt^ method by normalizing it to GAPDH expression. The activation sgRNAs used in this study and the qPCR primers were listed in Supplementary information Table S1.

### Activity and specificity assessment

For the comparison of the performance among the CRISPR systems with different promoters, Tag-seq results were used for calculating the activity and specificity. Activity value was calculated as the mean ratio of the on-target reads across all the tested sites, normalized to the CAG-driven CRISPR-Cas12a system. The specificity Index was calculated as the ratio of the on-target reads to the on-target reads plus the off-target reads across all the tested sites.

## Results

### Effects of promoters in transient transgene expression and activity of CRISPR-Cas12a editors

To explore the impact of promoters on gene expression and CRISPR editors, we chose four promoters (CAG,∼1700bp; EF1a core, ∼210bp; CMV, ∼500bp; and PGK, ∼500bp) for the investigations, because they were commonly used in the design of the CRISPR-Cas systems. And we focused on the CRISPR-Cas12a editor because it is also a versatile gene-editing tool and retains unique features distinguished from the widely-used CRISPR-Cas9, such as maintaining higher specificity and holding self-processing ability thus enabling to do multiplex editing with a single crRNA transcription[27-30]. We constructed the plasmids by employing the enAsCas12a-HF nuclease (an enhanced activity mutant of AsCas121[31]) that was driven by these four promoters and fused with a 3xHA tag and a P2A-mCherry reporter (Fig. 1a), as this variant could easier induce off-target editings[25, 31] and this property made it more suitable for doing specificity comparison. First, we detected the gene expression level in these four plasmids by transfecting them in MCF7 cells. Western bolting results showed that the CAG promoter-driving plasmid displayed the highest expression level of Cas12a-protein (anti-HA), followed by promoter CMV, EF1a, and PGK (Fig. 1b), and the detection of the mCherry reporter expression by FACS confirmed the similar results (Fig. 1c), both of which were consistent with the previous study that the CAG promoter maintained robust transgene expression in human cells[16].

**Fig. 1.**
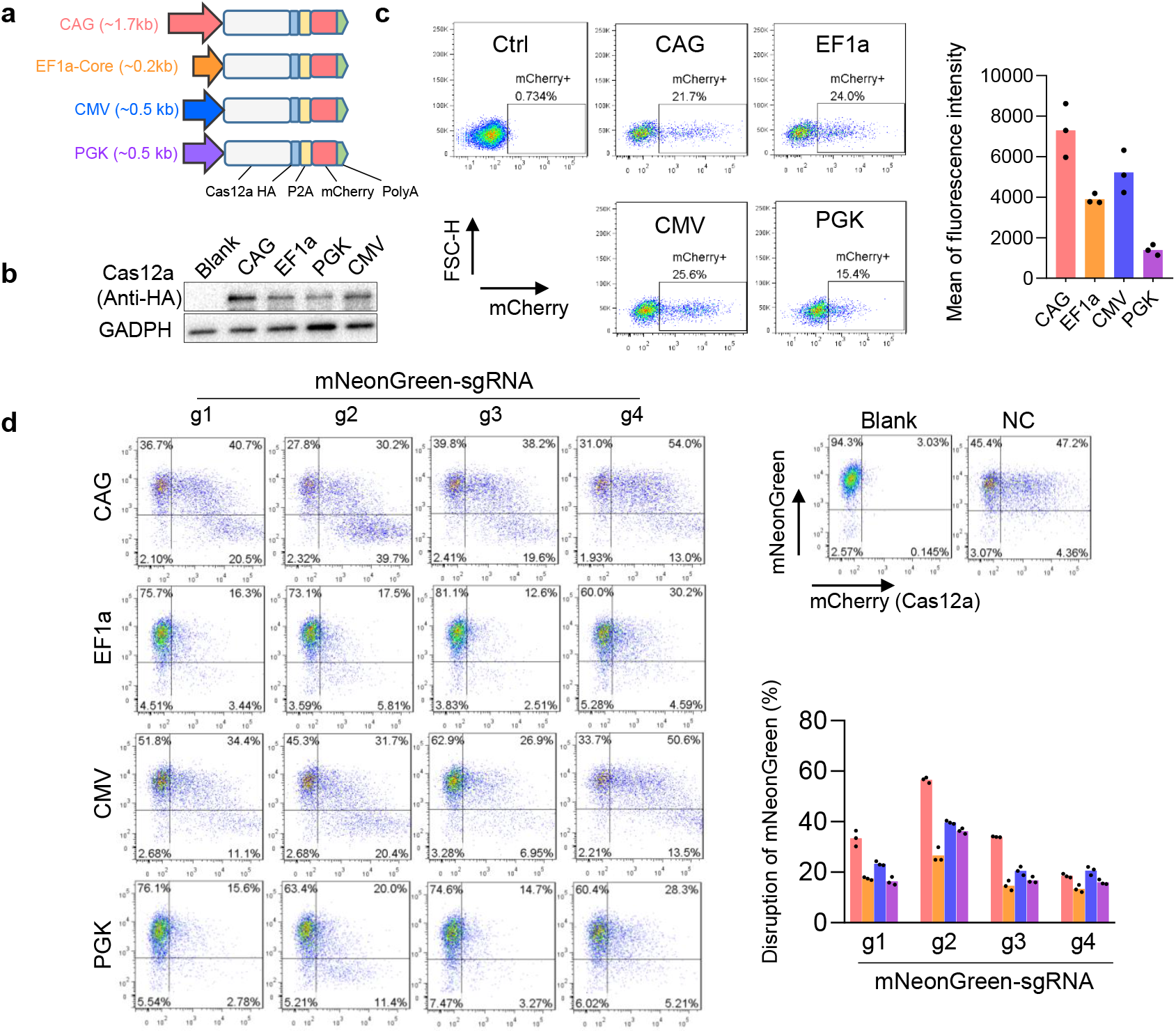
Comparison of the expression level and the activity of the CRISPR-Cas12a systems using different promoters. **a** Schematic of the CRISPR-Cas12a systems driven by different promoters (CAG, EF1a Core, CMV, and PGK). **b** Western blot showing the expression levels of the Cas-protein nucleases (anti-HA) driven by different promoters. Blank, HEK293T without transfection. **c** FACS detected the mCherry expression level driven by different promoters in MCF7 cells. Mean values are presented with SEM, n=3 independent experiments. **d** FACS analyses of the editing activities of the CRISPR-Cas12a systems with different promoters with four sgRNAs that targeted the mNeonGreen in the HEK293T KI mNeonGreen reported cell line. The editing efficiency was determined as the proportion of mNeonGreen negative cells within the Cas-nucleases transfected cells (mCherry-positive). sgRNA1/2/3/4, sgRNAs targeting mNeonGreen. Mean values are presented with SEM, n=3 independent experiments. Blank, cells without transfection. NC, cells transfected with a non-targeted sgRNA.

Since the promoters affected the Cas12a-nucleases expression, we then wondered about their impacts on cleavage activity in genome editing. We performed the tests by transfecting the four plasmids with the sgRNAs that targeted four loci of the mNeonGreen in the HEK293T-KI mNeonGreen reporter cells. As a result, the FACS data showed that the CAG promoter driving enAsCas12a-HF editor induced the most negative mNeonGreen in all the four tested sites, followed by promoter CMV, and then EF1a and PKG (Fig. 1d). Together, these data revealed that different promoters affected the cutting activities of the CRISPR editors.

### Effects of promoters in CRISPR-Cas12a systems on targeting specificity

As a stronger promoter boosted the higher cleavage activity of the editor (Fig. 1d) and the off-target effect was a key concern of the CRISPR tools for therapeutic applications, we next wondered whether they also affected the targeting accuracy of the CRISPR-Cas12a systems. To this end, after detection of their transfection efficiency (Supplementary Fig. 1), we performed Tag-seq assays[24] in MCF7 cells with 25 sgRNAs targeted 17 genes (Fig. 2a, b and Supplementary Fig. 2) to compare their genome editing specificities, because the Tag-seq method enables to parallelly profile the off-target cleavages induced by Cas-protein at diverse sites in a single transfection, which is a rapid and cost-efficient approach for evaluating the performance of a nuclease[24]. Consistent with Fig. 1d, Tag-seq results showed that the CAG promoter driving enAsCas12a-HF editor had the highest level of editing efficiency, followed by promoter CMV with ∼92% (vs CAG), and then EF1a with ∼61%(vs CAG) and PGK with ∼59% (vs CAG) (Fig. 2c), however, the specificities did not markedly affect (Fig. 2d, e). Exception for MCF7 cells, we also performed the experiments in HEK293T cells and similar results were obtained (Supplementary Fig. 3). Together, these data indicated that the promoters affected the activity of the Cas12a editor but with little impact on targeting specificity.

**Fig. 2.**
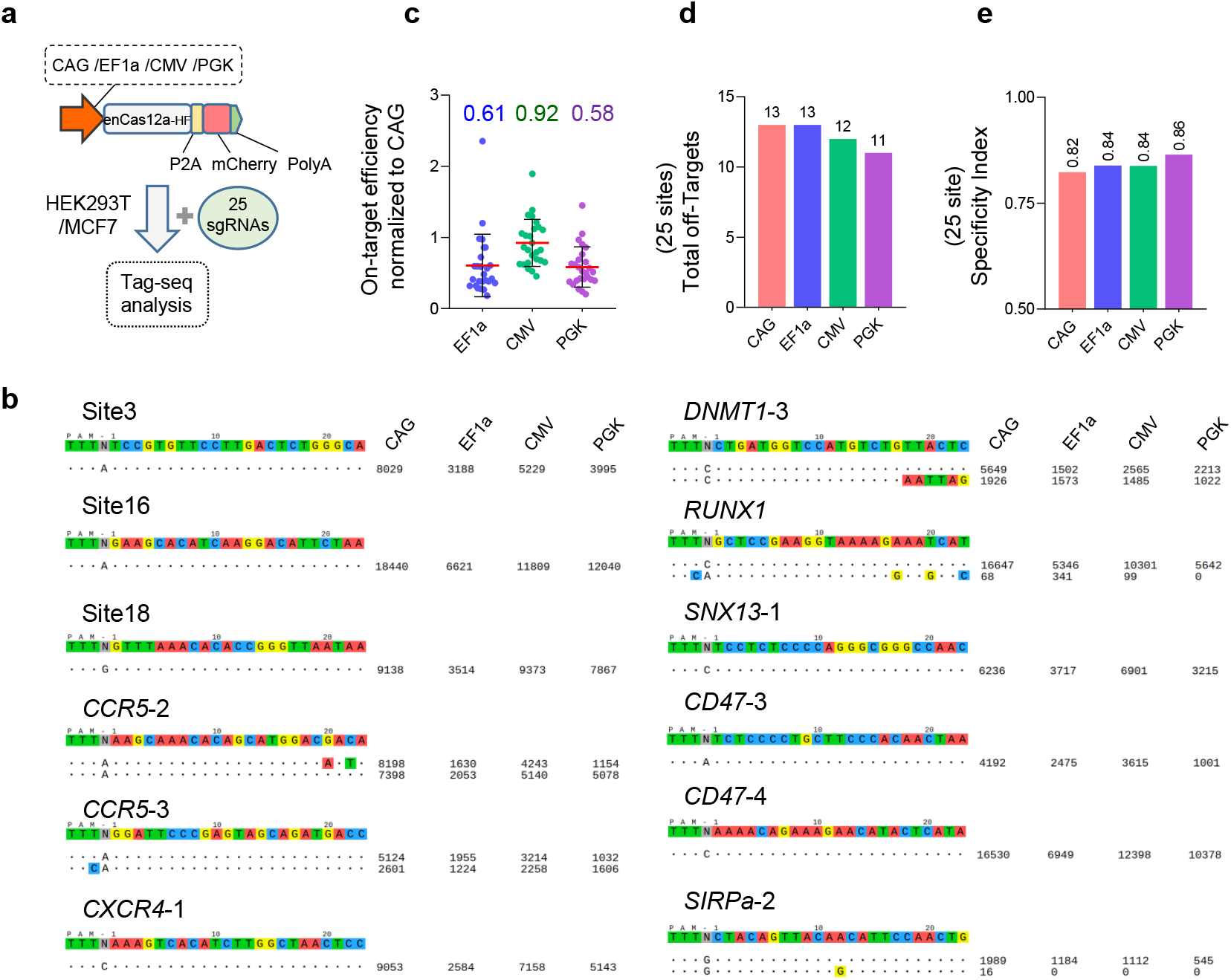
Editing specificity comparison of the CRISPR-Cas12a systems with different promoters. **a** Schematic of the editing specificity analysis by Tag-seq among the CRISPR-Cas12a systems with different promoters. **b** Tag-seq-based comparative analysis of CRISPR-Cas12a systems with different promoters (also see Supplementary Fig. S2). For visualization, the sgRNA sequences were shown at the top, and the on-target and off-target sites were shown without or with mismatches to the sgRNA sequence by color highlighting. Sequencing read counts were shown to the right of each site. **c** Normalization of on-target activity of the various CRISPR-Cas12f1 systems to the CRISPR-Cas12f1 driven by the CAG promoter, value= (other systems on-target reads)/(CRISPR-Cas12f1 with CAG promoter). **d** Total number of off-target sites detected with the twenty-five sgRNAs. **e** Specificity Index assessment (value was calculated by the ratio of total on-target reads to the on-target reads plus the off-target reads within the twenty-five sites).

### Effects of promoters in CRISPR-Cas12a systems on multiplex editing

As mentioned above, one unique feature of the Cas12a enzyme over the widely-used Cas9 is the multiplex editing, where Cas12a can process multiple functional crRNAs from a single long transcription to simplify multiplex targeting in cells and animals[29, 30, 32], and this feature also makes the Cas12a as a powerful approach for versatile gene modulation with applications in cell reprogramming and combinatorial genetic screening[33, 34]. To assess the impact of the promoters on this property of the Cas12a nuclease, we cloned a long crRNA transcription targeting six sites, including *DNMT1, EMX1, CTLA4, CCR5, SIPRa*, and *RUNX1* (Fig. 3a). Agreed with the above results, Tag-seq assays showed that the Cas12a editors with different promoters could mediate these six sites editing (Fig. 3b), but with various levels, where the CAG promoter driving enAsCas12a-HF nuclease displayed the highest efficiency, followed by promoter CMV, and then EF1a and PKG (Fig.3c). However, for the specificity, the CAG and the CMV promoters driving effectors exhibited relatively lower than that of EF1a and PGK (Fig.3d, e), indicating a slight impact on specificity. And similar results could be observed in the HEK293T cells (Supplementary Fig. 4). Again, these data demonstrated that with a sight compromise in specificity, promoters robustly affected the activity of the Cas12a editor in multiplex editing.

**Fig. 3.**
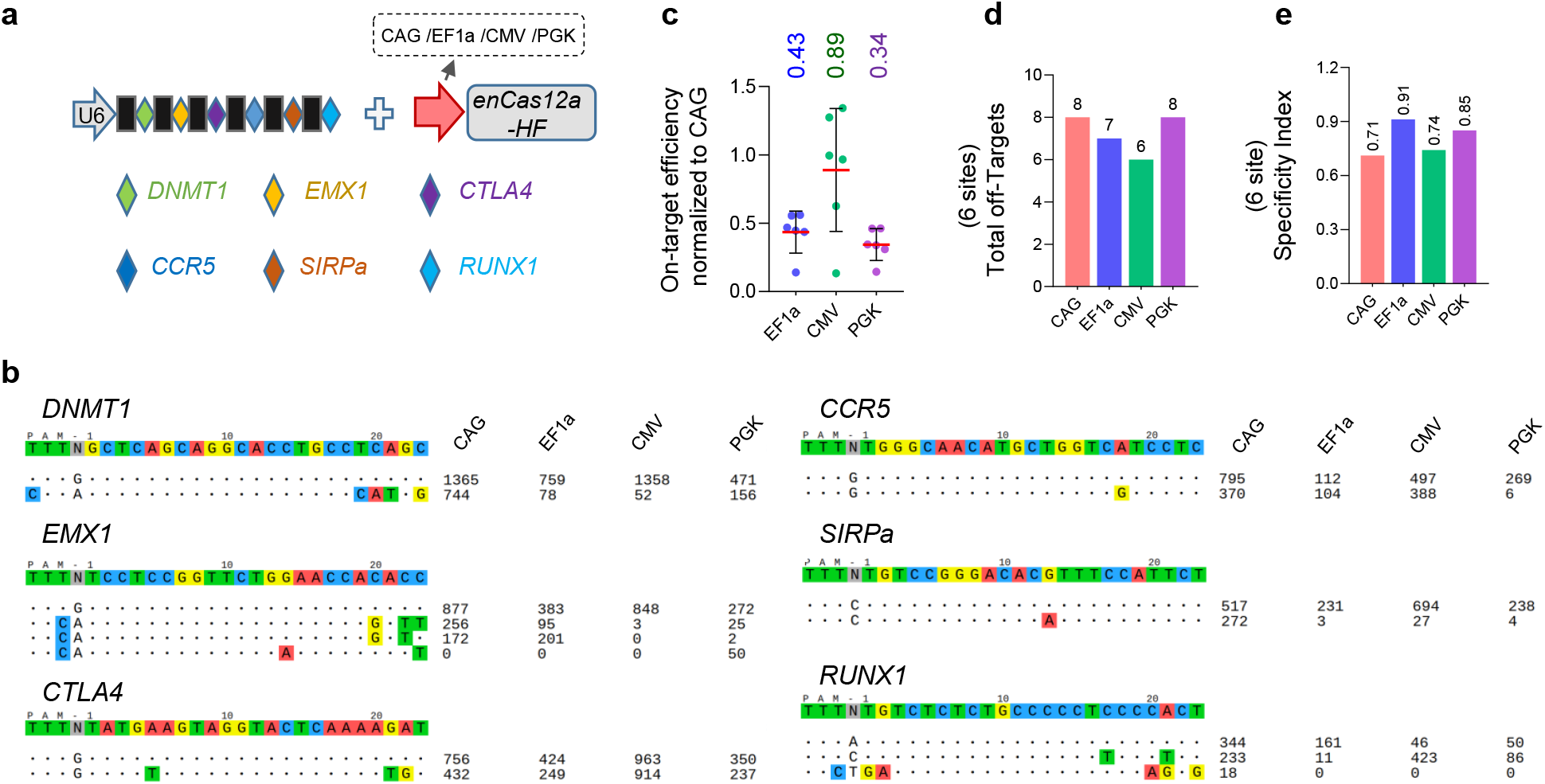
Multiplex-editing specificity comparison of the CRISPR-Cas12a systems with different promoters. **a** Schematic of the multiplex editing. **b** Tag-seq-based comparative analysis of CRISPR-Cas12a systems with different promoters in multiplex editing. **c** Normalization of on-target activity of the various CRISPR-Cas12f1 systems to the CRISPR-Cas12f1 driven by the CAG promoter in multiplex editing, value= (other system on-target reads)/(CRISPR-Cas12f1 with CAG promoter). **d** Total number of off-target sites detected with the six sgRNAs in multiplex editing. **e** Specificity Index assessment (value was calculated by the ratio of total on-target reads to the on-target reads plus the off-target reads within the six sites).

### Effects of promoters in CRISPR-Cas12a systems on gene activation

CRISPR-based activation (CRISPRa) activator is also a promising gene-editing tool of the CRISPR-Cas system and has been proven to have great potential in therapy applications[3-5]. Next, we tested the effect of the promoters on this gene-editing technique. We constructed the CAG/EF1a/CMV/PGK promoter driving enAsCas12a-HF-based activators by fusing the DNase-inactive enAsCas12a-HF to the synthetic VPR (VP64-p65-Rta) activation domain (Fig. 4a) and detected their transcriptional activation of *MYOD, IL1RN*, and *HBG* in MCF7 cells. As a result, we found that similar to the genome-cleaving editors, the activator with a CAG promoter maintained a higher activation level than that of other systems (Fig. 4b-d). As the same, we performed the experiments in HEK293T cells and a similar conclusion was obtained (Supplementary Fig. 5). These data suggested that the promoters could also affect the CRISPR-Cas12a system for gene activation.

**Fig 4.**
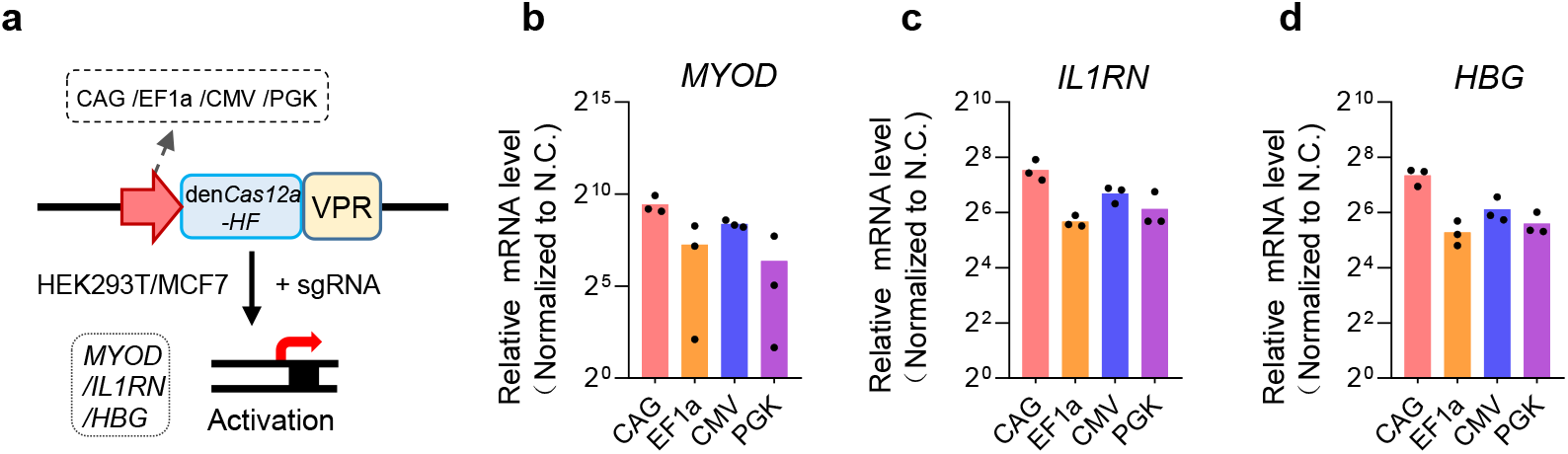
Transcriptional activation ability comparison of the CRISPR-Cas12a systems with different promoters. **a** Schematic of the gene activation systems based on CRISPR-Cas12a systems with different promoters. VPR, synthetic VP64-p65-Rta activation domain. **b-d** qPCR analysis of the transcriptional activation among the CRISPR-Cas12a systems with different promoters guided by a single sgRNA targeting each promoter region of *MYOD* (b), *IL1RN* (c), and *HBG* (d) in human MCF7 cells. Mean values are presented with SEM, n=3 independent experiments.

### Effects of promoters in CRISPR-Cas12a systems on base editing

Another powerful application of the CRISPR-Cas system is the base editor, which allows making the single base change in the genome without inducing double-stranded DNA breaks or donor DNA templates, and without reliance on homology-directed repair (HDR) [8, 9], and it has been demonstrated to be a promising technology for basic research and translational medicine therapy[10, 13]. Finally, we profiled the impact of the promoters on the Cas12a base editor.

We chose Site 3 and Site 16 as the tests because these two sites had been well-studied in previous studies[31, 35]. As the above results (Fig. 1-4) showed that the PGK promoter displayed a similar performance to the EF1a promoter, we thus only chose the CAG, EF1a, and CMV for further investigations. We designed CAG/EF1a/CMV driving enAsCas12a-HF-based Cytosine base editors (CBE) editors by fusing the DNase-inactive enAsCas12a-HF to the rat APOBEC deaminase (Fig. 5a) because this system had been reported significantly improved the base editing ability of the AsCas12a nuclease[31]. We first determined the base editing ability by targeting Site3 in MCF7 cells and sanger sequencing revealed that the C9, C10, C15, and C17 of Site 3 had a significant double peak phenomenon (Fig. 5b), suggesting that these sites were edited by the enAsCas12a-HF-based CBE editors. Next, we performed the Deep-seq assay for a more detailed comparison of these systems. As a result, Deep-seq data showed that the CAG promoter driving editor was the most active, followed by the CMV (Fig. 5c, d), and then the EF1a, and the tests were administrated in another locus Site16, which exhibited a similar result (Supplementary Fig. 6a, b). Next, to test the universality of this conclusion, we also performed similar experiments in the HEK293-T cell line and the same data were obtained (Supplementary Fig. 6c, f). All these results revealed that the promoters had an impact on the base editing ability of the CRISPR-Cas12a system as well.

**Fig 5.**
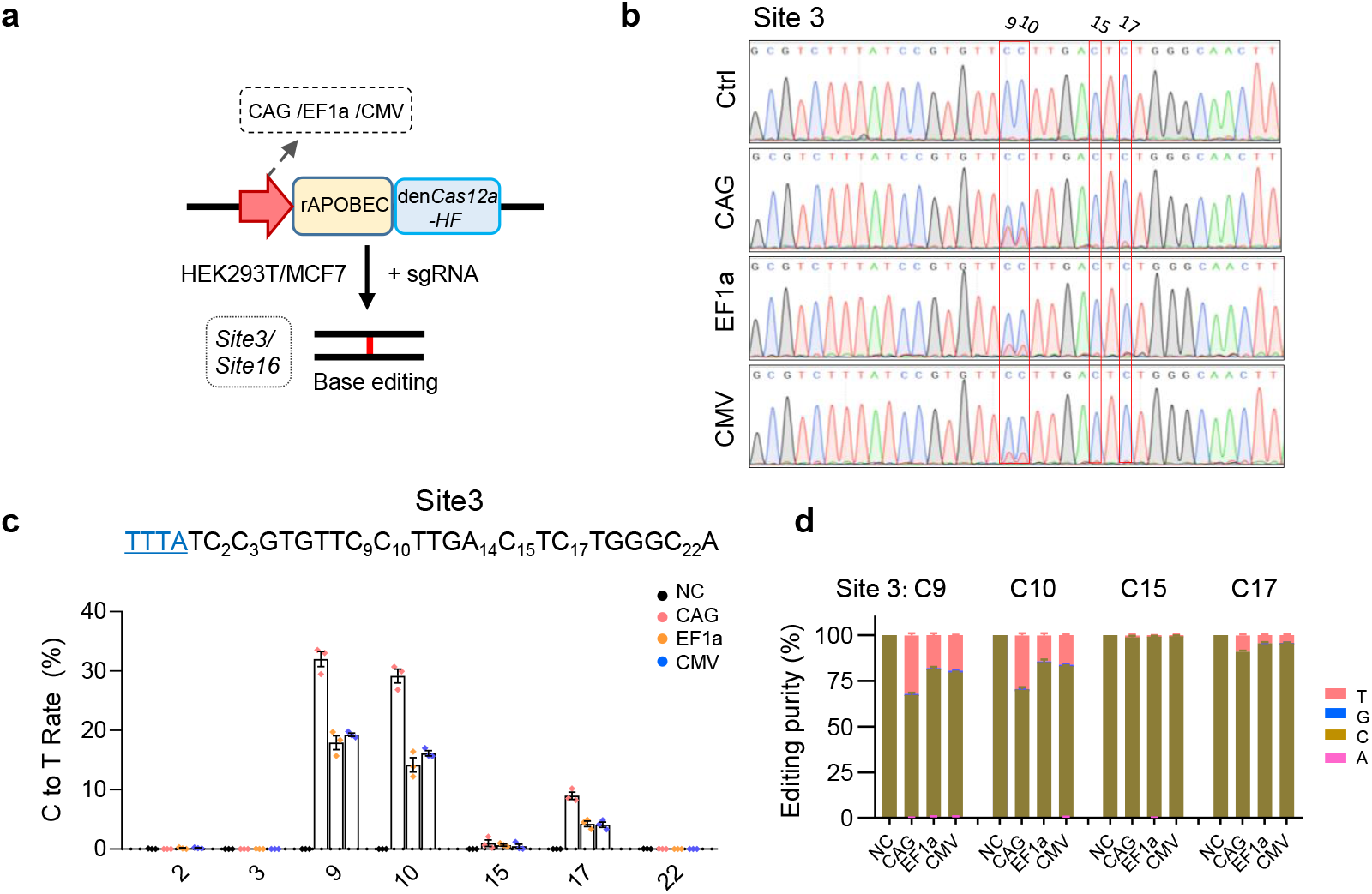
Base editing comparison of the CRISPR-Cas12a systems with different promoters. **a** Schematic of the base editing systems based on CRISPR-Cas12a with different promoters. rAPOBEC1, rat APOBEC1. **b** Sanger sequencing analysis of the base editing ability at site 3. **c** Deep-seq revealed the cytosine to thymine (C-to-T) editing at site 3. Mean values are presented with SEM, n=3 independent experiments. **d** Analysis of the editing purity at site 3. The fraction was plotted by calculating each nucleotide read within the total reads at this site. Mean values are presented with SEM, n=3 independent experiments. NC, cells transfected with a non-targeted sgRNA.

## Discussion

A CRISPR editor with a distinct purpose could get better performance from the rational design of its Cas-protein expression cassette. As our data showed, the CRISPR-Cas12a tools with different promoters exhibited various editing performances, we thus highly recommended using these promoters according to the fitness of their unique features to the intended research. For instance, consistent with the previous report[16], our data also revealed that the Cas12a driven by the CAG promoter maintained a high-level expression in the human cell line and thus boosting the editing activities with a comparable targeting accuracy (Fig.1, 2), however, the size of the CAG promoter was relatively large (∼1700 bp, Table 1). Therefore, we would recommend using this promoter for a purpose that requires a robust activity but has no size limitation, such as disrupting genes by transient transfection in cell lines or *in-vitro* applications because the editing activity is generally a priority in these experiments. On the contrary, for the investigations related to *in vivo* therapy applications, we would recommend using the EF1a core promoter, since this promoter was much smaller with only ∼210 bp, which could be more easily packaged into AAV vectors for *in vivo* delivery. What’s more, it maintained a higher targeting accuracy, and the specificity (and thus the safety) was always a priority in the therapeutic applications. For more detailed comparisons of the performance of the promoters in CRISPR-Cas systems see Table 1.

**Table 1.** Comparison of promoters in CRISPR-Cas12a editors

| **Promoter**<br>**Contents** | **CAG** | **EF1a Core** | **CMV** | **PGK** |
| --- | --- | --- | --- | --- |
| Size | ~1700 bp | ~210 bp | ~500 bp | ~500 bp |
| GC content | ~66 % | ~59 % | ~49 % | ~65 % |
| Activity* | 100% | 40~60% | 70~90% | 40~60% |
| Specificity* | ~75% | ~84% | ~78% | ~82% |
| Off-target No.<br>(in 25 sgRNAs ) | 19 | 14 | 18 | 14 |
| Application<br>scenarios | Need robust activity<br>without size limitation | Require a compact<br>size design | An alternative for<br>CAG | Require a lower<br>lever of Cas-protein |

Note: \*Activity was calculated by normalizing the efficiency to the CAG-driven Cas12a editor. \*Specificity index was calculated by Tag-seq results as the ratio of the on-target reads to the on-target reads plus the off-target reads across all the tested sites in this study.

## Conclusion

In summary, the comparison outlines the editing performance of four commonly used promoters in the CRISPR-Cas12a system, which provides the basic information for the gene-editing toolkit and can be a guideline, as well as a valuable resource for this field.

## Abbreviations

AAV: adeno-associated virus
VPR: VP64-p65-Rta
PEI: polyethyleneimine
enAsCas12a: activity enhanced AsCas12a
CRISPRa: CRISPR-based activation
CBE: Cytosine base editors
HDR: homology-directed repair

## Declarations

### Acknowledgments

We are grateful to all members for the helpful comments and discussions on the manuscript.

### Author Contributions

H.H. and M.Z. conceived the project, designed the experiments, and wrote the manuscript. J.L. and Q.L. performed most of the experiments and conducted the data analysis. H.Z., help to analyze the data and edit the manuscript. All authors had read and approved the final manuscript.

### Data availability

The sequencing data related to this study have been deposited in NCBI (Bioproject PRJNA884019). And other data that support this study are available from the corresponding author upon reasonable request.

### Ethics approval and consent to participate

Not applicable.

---

**Figure S1.**
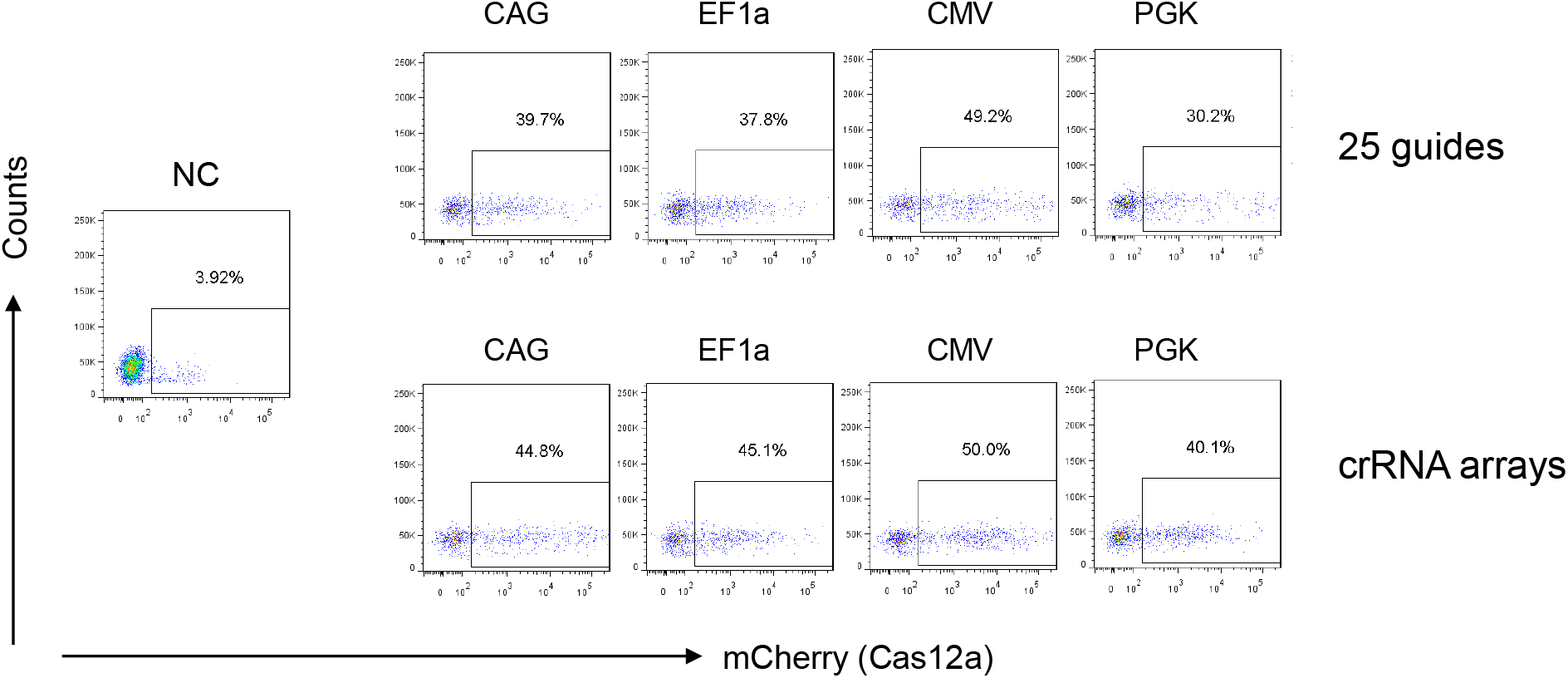
Detection of the transfection efficiency by FACS. The transfection were administrated by PEI-based method in 12-well plates with the plasmids expression of Cas-protein (fusing a P2A mCherry reporter, 600 ng) and the pooled sgRNAs (up, 25 guides, total 600 ng) or the crRNA array (down, 6 sites, 600 ng), and an Tag-oligo DNA (10uM), and the transfection efficiency was determined by FACS with the mCherry reporter.

**Figure S2.**
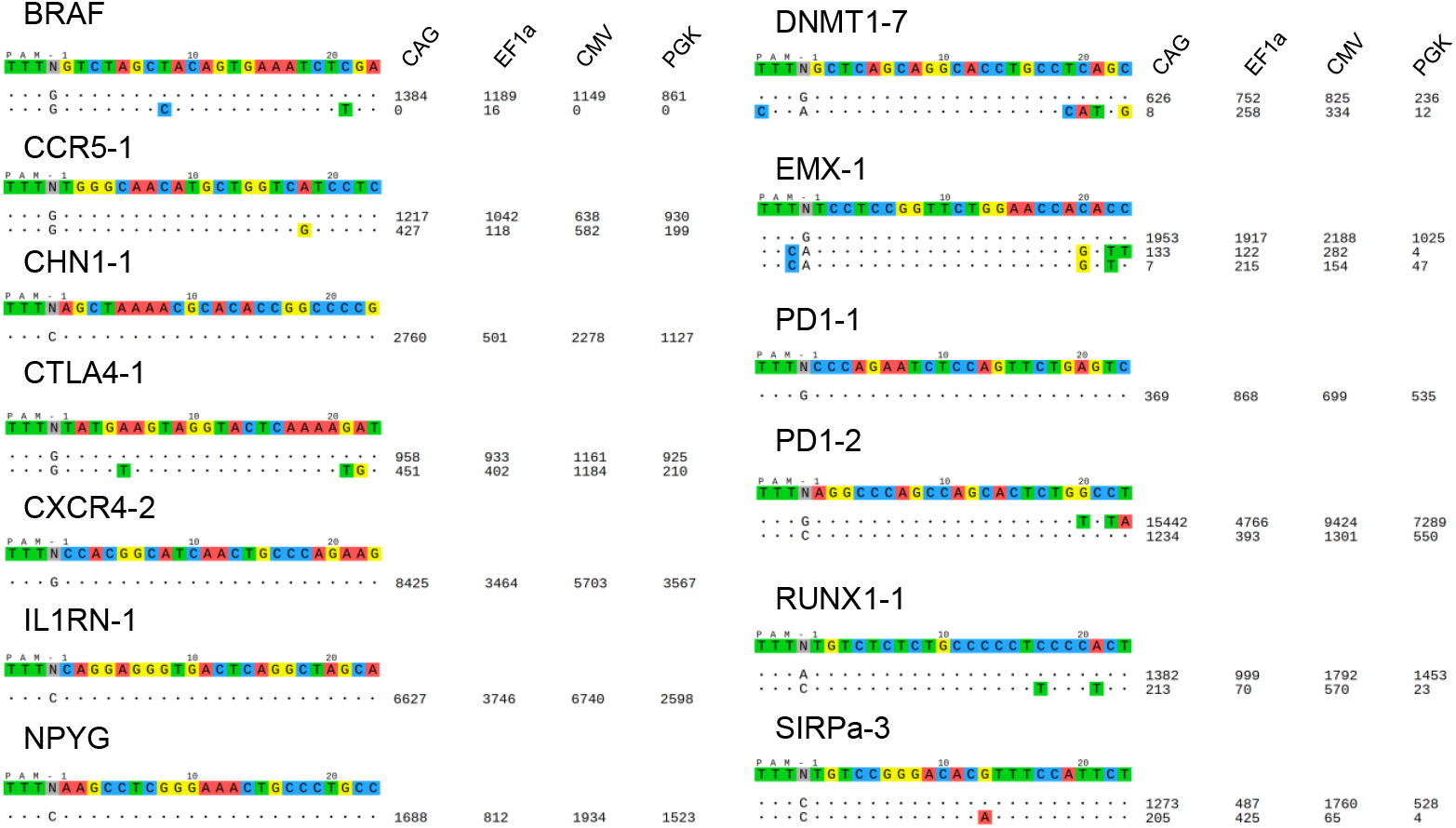
Detection of the editing specificity by Tag-seq. MCF7 cells were co-transfected with the plasmids expressing AsCas12a-HF driven by different promoters, a pooled twenty-five sgRNAs, and the donor Tag sequence. Genomic DNA was harvested after three days post-transfection for libraries construction and Tag-seq analysis. Read counts represented a measure of cleavage frequency at a given site, mismatched positions showing the off-targets were highlighted in color within the spacer or PAM. (also see Fig. 2b)

**Figure S3.**
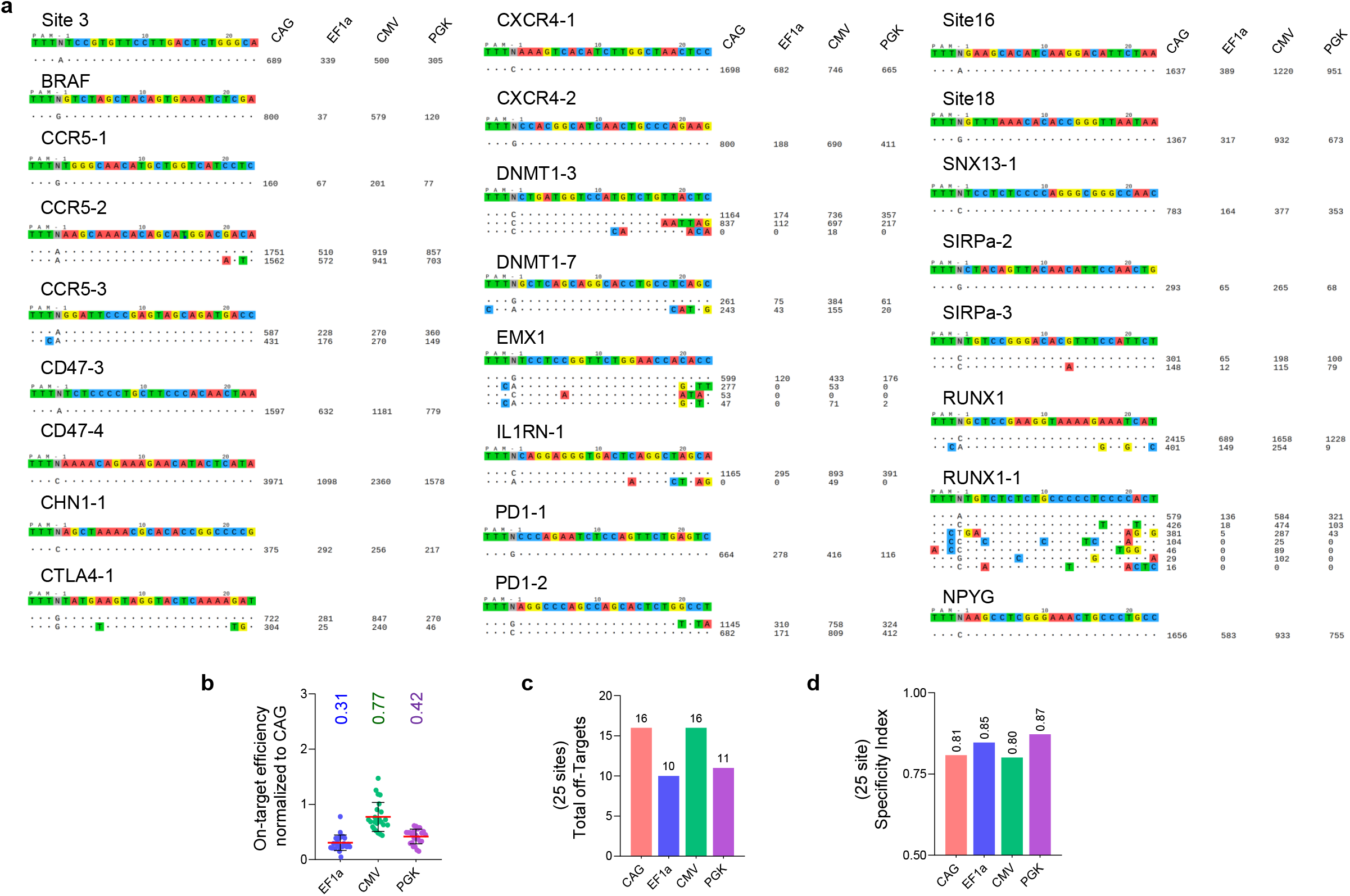
The specificity of the CRISPR-Cas12a systems using different promoters in HEK293T cells. **a** Detection of the editing specificity by Tag-seq in HEK293T cells. HEK293T cells were co-transfected with the plasmids expressing AsCas12a-HF driven by different promoters, a pooled twenty-five sgRNAs, and the donor Tag sequence. Read counts represented a measure of cleavage frequency at a given site, mismatched positions showing the off-targets were highlighted in color within the spacer or PAM. **b** Normalization of on-target activity of the various CRISPR-Cas12a systems to the CRISPR-Cas12a driven by the CAG promoter, value=(other systems on-target reads)/(CRISPR-Cas12a with CAG promoter). **c** Total number of off-target sites detected with the twenty-five sgRNAs. **d** Specificity Index assessment (value was calculated by the ratio of total on-target reads to the on-target reads plus the off-target reads within the twenty-five sites).

**Figure S4.**
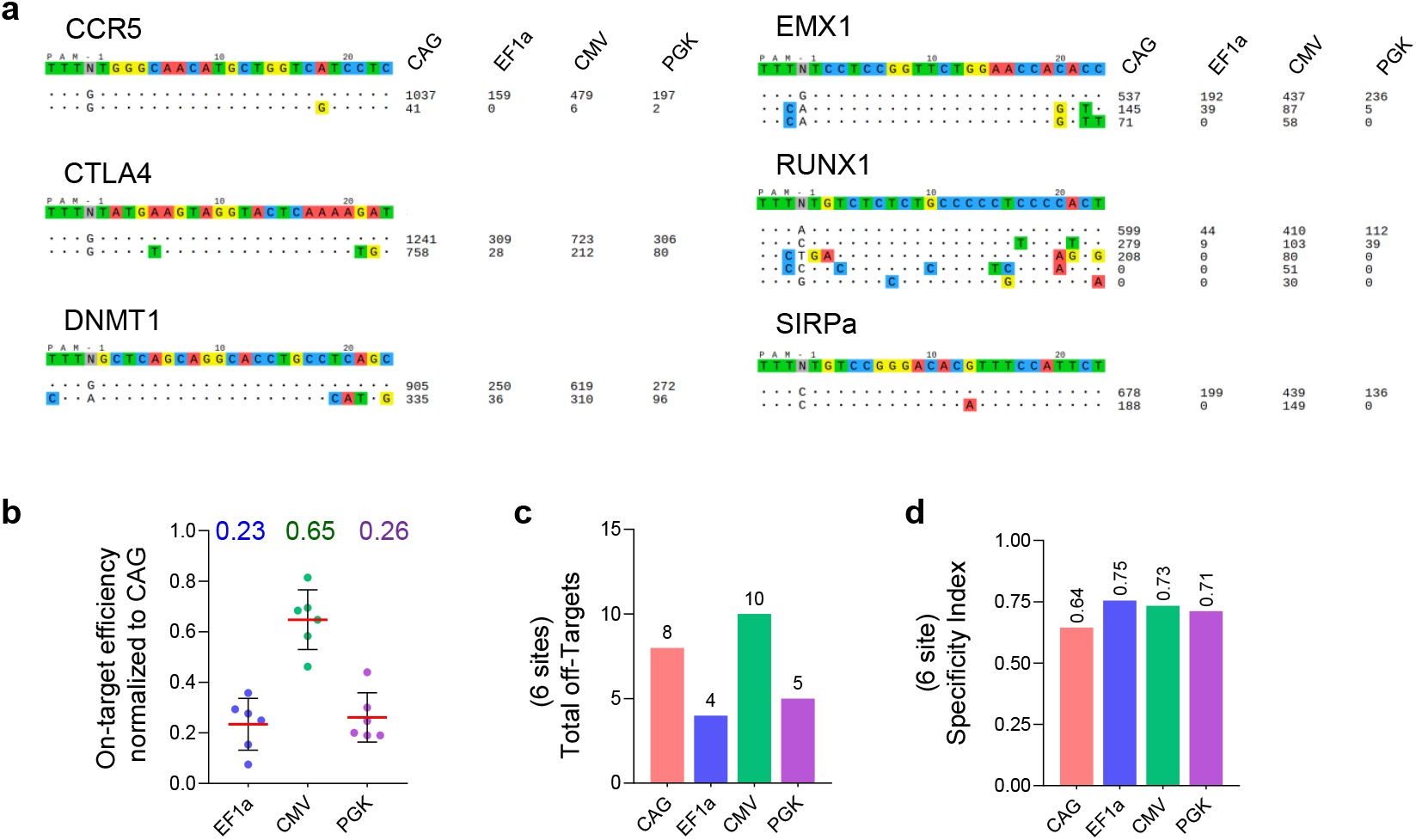
The multiplex-editing specificity of the CRISPR-Cas12a systems with different promoters in HEK293T cells. **a** Detection of the multiplex-editing specificity by Tag-seq in HEK293T cells. HEK293T cells were co-transfected with the plasmids expressing AsCas12a-HF driven by different promoters, a crRNA array targeting six sites, and the donor Tag sequence. Read counts represented a measure of cleavage frequency at a given site, mismatched positions showing the off-targets were highlighted in color within the spacer or PAM. **b** Normalization of on-target activity of the various CRISPR-Cas12a systems to the CRISPR-Cas12a driven by the CAG promoter, value=(other systems on-target reads)/(CRISPR-Cas12a with CAG promoter). **c** Total number of off-target sites detected with the six sites. **d** Specificity Index assessment (value was calculated by the ratio of total on-target reads to the on-target reads plus the off-target reads within the six sites).

**Figure S5.**
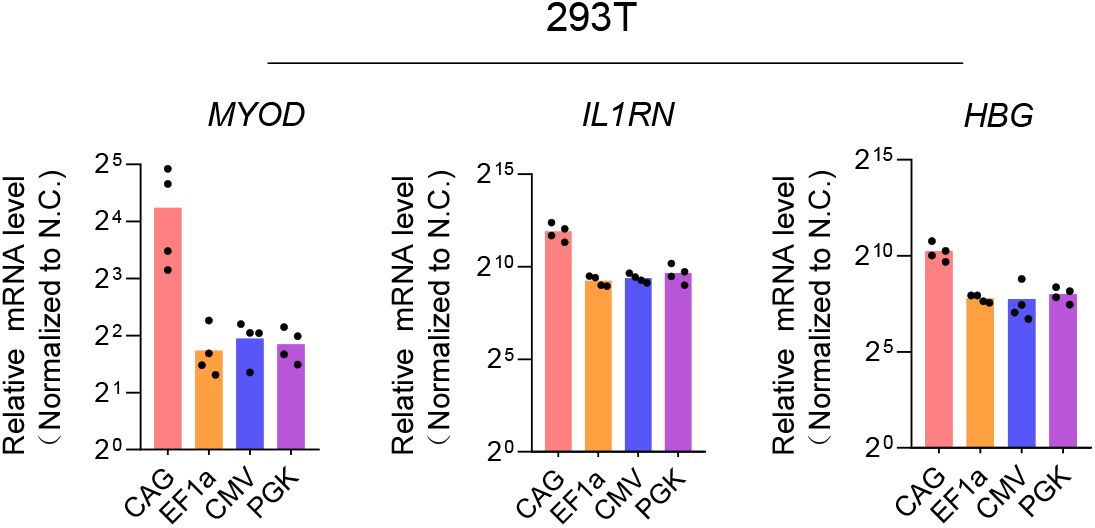
The activation level of the Cas12a-based CRISPRa systems using different promoters in HEK293T cells. **a-c** qPCR analysis of the transcriptional activation among the CRISPR-Cas12a systems with different promoters guided by a single sgRNA targeting each promoter region of *MYOD* (a), *IL1RN* (b) and *HBG* (c) in human HEK293T cells. Mean values are presented with SEM, n=4 independent experiments.

**Figure S6.**
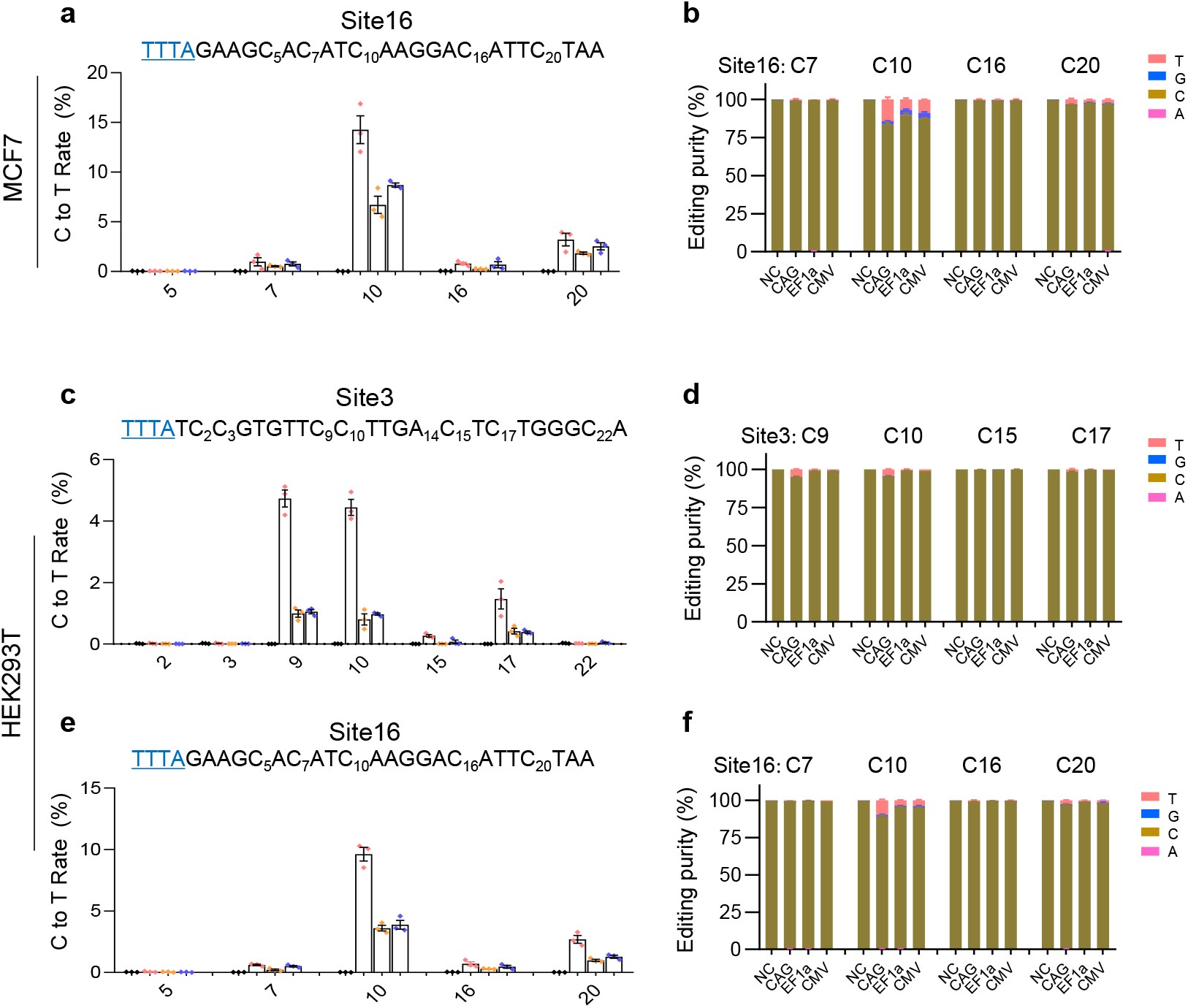
Base editing activity of the CRISPR-Cas12a systems with different promoters. **a** Deep-seq revealed the cytosine to thymine (C-to-T) editing at site 16 in MCF7 cells. **b** Analysis of the editing purity at site 16. The fraction was plotted by calculated each nucleotide reads within total reads at this site. **c** Deep-seq revealed the cytosine to thymine (C-to-T) editing at site 3 in HEK293T cells. **d** Analysis of the editing purity at site 3. **e** Deep-seq revealed the cytosine to thymine (C-to-T) editing at site 16 in HEK293T cells. **f** Analysis of the editing purity at site 16. For all the figures, mean values are presented with SEM, n=3 independent experiments. NC, cells transfected with a non-targeted sgRNA.

## Supplementary Information

**Table S1.**
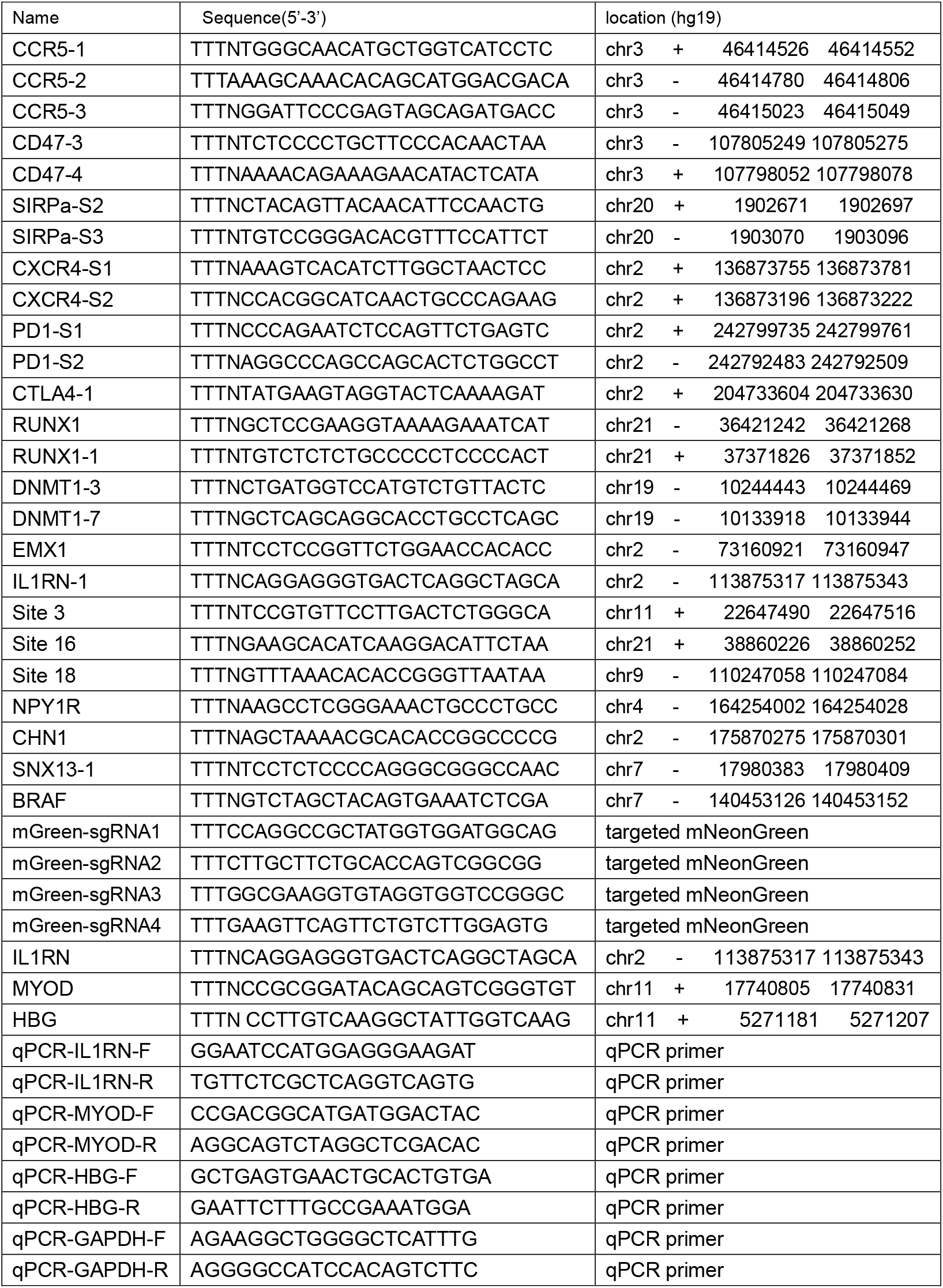
The sgRNAs and primers used in this study

**Table S2.**
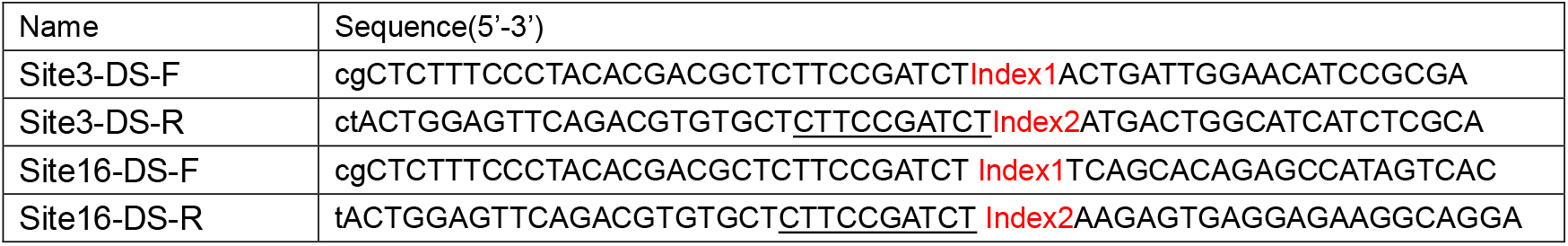
Deep-seq primers for this study

### DNA sequences

#### The CAG promoter (1719 bp)

gacattgattattgactagttattaatagtaatcaattacggggtcattagttcatagcccatatatggagttccgcgttacataacttacggtaaatggcccgcctgg

ctgaccgcccaacgacccccgcccattgacgtcaataatgacgtatgttcccatagtaacgccaatagggactttccattgacgtcaatgggtggactatttacg

gtaaactgcccacttggcagtacatcaagtgtatcatatgccaagtacgccccctattgacgtcaatgacggtaaatggcccgcctggcattatgcccagtacat

gaccttatgggactttcctacttggcagtacatctacgtattagtcatcgctattaccatgggtcgaggtgagccccacgttctgcttcactctccccatctccccccc

ctccccacccccaattttgtatttatttattttttaattattttgtgcagcgatgggggcggggggggggggggcgcgcgccaggcggggcggggcggggcgag

gggcggggcggggcgaggcggagaggtgcggcggcagccaatcagagcggcgcgctccgaaagtttccttttatggcgaggcggcggcggcggcggc

cctataaaaagcgaagcgcgcggcgggcgggagtcgctgcgttgccttcgccccgtgccccgctccgcgccgcctcgcgccgcccgccccggctctgactg

accgcgttactcccacaggtgagcgggcgggacggcccttctcctccgggctgtaattagcgcttggtttaatgacggctcgtttcttttctgtggctgcgtgaaag

ccttaaagggctccgggagggccctttgtgcgggggggagcggctcggggggtgcgtgcgtgtgtgtgtgcgtggggagcgccgcgtgcggcccgcgctgc

ccggcggctgtgagcgctgcgggcgcggcgcggggctttgtgcgctccgcgtgtgcgcgaggggagcgcggccgggggcggtgccccgcggtgcgggg

gggctgcgaggggaacaaaggctgcgtgcggggtgtgtgcgtgggggggtgagcagggggtgtgggcgcggcggtcgggctgtaacccccccctgcacc

cccctccccgagttgctgagcacggcccggcttcgggtgcggggctccgtacggggcgtggcgcggggctcgccgtgccgggcggggggtggcggcagg

tgggggtgccgggcggggcggggccgcctcgggccggggagggctcgggggaggggcgcggcggcccccggagcgccggcggctgtcgaggcgcg

gcgagccgcagccattgccttttatggtaatcgtgcgagagggcgcagggacttcctttgtcccaaatctgtgcggagccgaaatctgggaggcgccgccgca

ccccctctagcgggcgcggggcgaagcggtgcggcgccggcaggaaggaaatgggcggggagggccttcgtgcgtcgccgcgccgccgtccccttctcc

atctccagcctcggggctgtccgcagggggacggctgccttcgggggggacggggcagggcggggttcggcttctggcgtgtgaccggcggctctagtgcct

ctgctaaccatgttcatgccttcttctttttcctacagctcctgggcaacgtgctggttattgtgctgtctcatcattttggcaaa

#### The EF1a core promoter (212 bp)

gggcagagcgcacatcgcccacagtccccgagaagttggggggaggggtcggcaattgatccggtgcctagagaaggtggcgcggggtaaactgggaa

agtgatgtcgtgtactggctccgcctttttcccgagggtgggggagaaccgtatataagtgcagtagtcgccgtgaacgttctttttcgcaacgggtttgccgccagaacacag

#### The CMV promoter (508 bp)

cgttacataacttacggtaaatggcccgcctggctgaccgcccaacgacccccgcccattgacgtcaataatgacgtatgttcccatagtaacgccaataggg

actttccattgacgtcaatgggtggagtatttacggtaaactgcccacttggcagtacatcaagtgtatcatatgccaagtacgccccctattgacgtcaatgacg

gtaaatggcccgcctggcattatgcccagtacatgaccttatgggactttcctacttggcagtacatctacgtattagtcatcgctattaccatggtgatgcggttttg

gcagtacatcaatgggcgtggatagcggtttgactcacggggatttccaagtctccaccccattgacgtcaatgggagtttgttttggcaccaaaatcaacggga

ctttccaaaatgtcgtaacaactccgccccattgacgcaaatgggcggtaggcgtgtacggtgggaggtctatataagcagagct

#### The PGK promoter (500 bp)

gggtaggggaggcgcttttcccaaggcagtctggagcatgcgctttagcagccccgctgggcacttggcgctacacaagtggcctctggcctcgcacacattc

cacatccaccggtaggcgccaaccggctccgttctttggtggccccttcgcgccaccttctactcctcccctagtcaggaagttcccccccgccccgcagctcgc

gtcgtgcaggacgtgacaaatggaagtagcacgtctcactagtctcgtgcagatggacagcaccgctgagcaatggaagcgggtaggcctttggggcagc

ggccaatagcagctttgctccttcgctttctgggctcagAGGCTGGGAAGGGGTGGGTCCGGGGGCGGGCTCAGGGGCGGGC

TCAGgggcggggcgggcgcccgaaggtcctccggaggcccggcattctgcacgcttcaaaagcgcacgtctgccgcgctgttctcctcttcctcatctccg

ggcctttcg

#### Expression of crRNA array

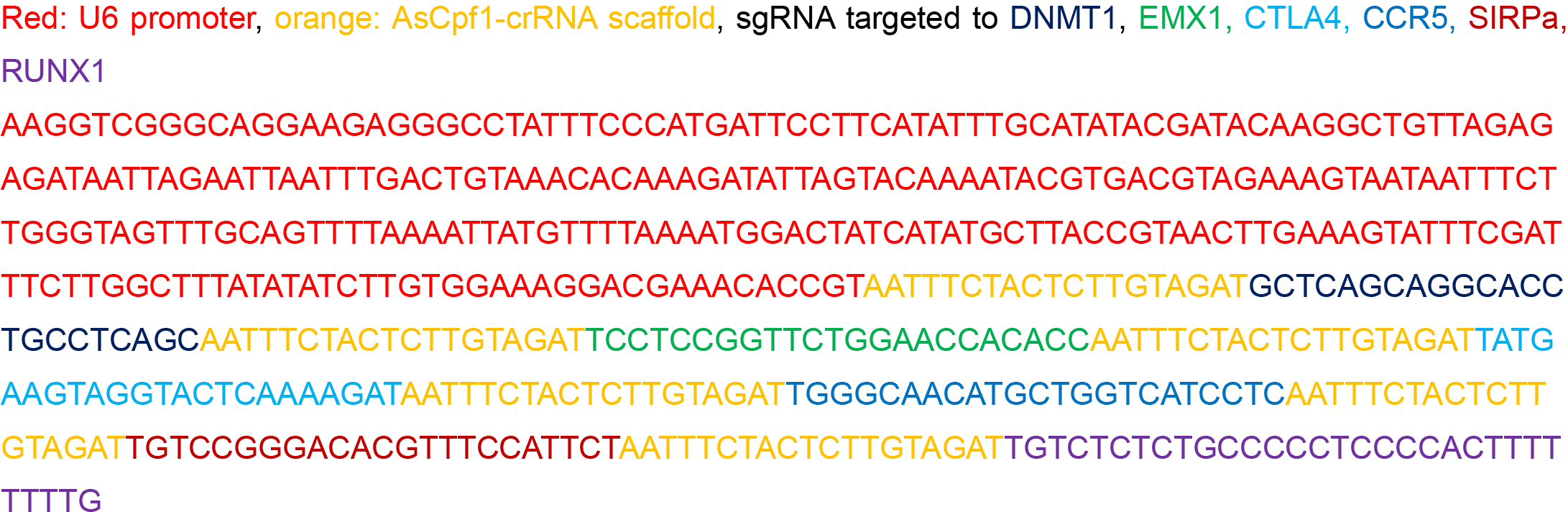

#### The amplicon sequence for Site 16

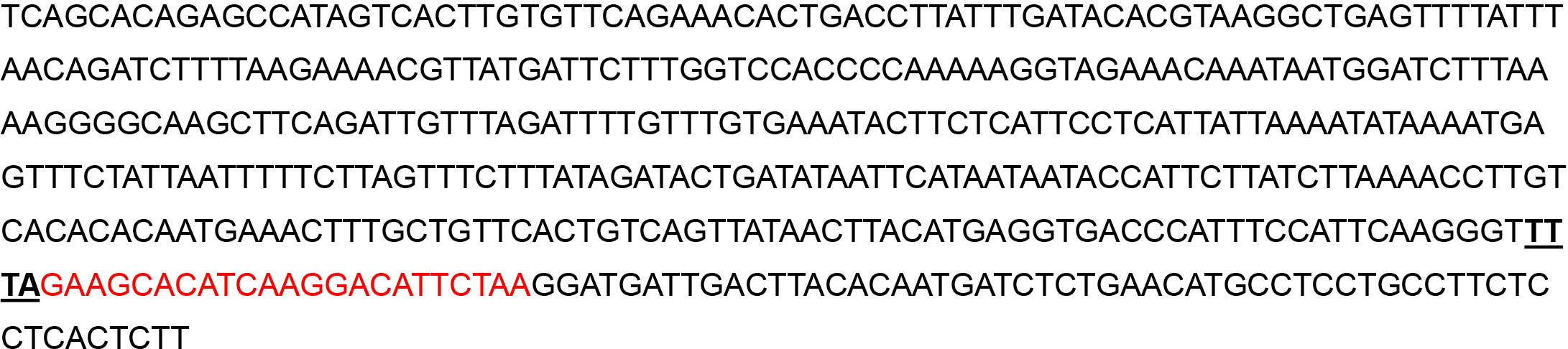

#### The amplicon sequence for Site 3

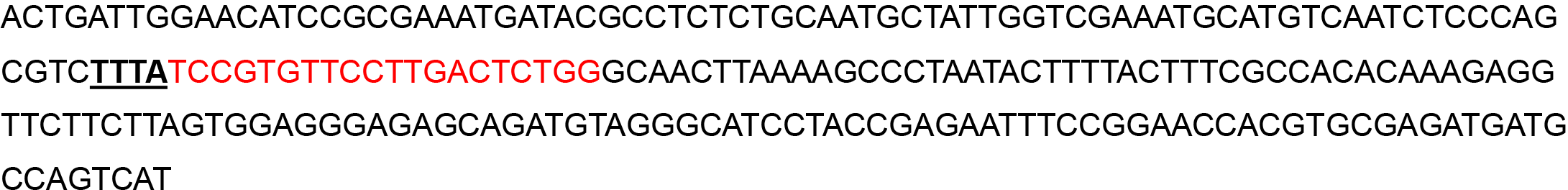

